# Broadband electronic resonance coherent anti-Stokes/Stokes Raman scattering microscopy

**DOI:** 10.1101/2025.09.23.677457

**Authors:** Yusuke Murakami, Norihide Sagami, Minori Masaki, Ryosuke Oketani, Masashi Yanagisawa, Hideaki Kano, Sakiko Honjoh, Kotaro Hiramatsu

**Affiliations:** International Institute for Integrative Sleep Medicine (WPI-IIIS), Tsukuba Institute for Advanced Research (TIAR), University of Tsukuba, 1-1-1 Tennodai, Tsukuba, Ibaraki 305-8575, Japan; Ph.D. Program in Humanics, University of Tsukuba, 1-1-1 Tennodai, Tsukuba, Ibaraki 305-8577, Japan; Department of Chemistry, Faculty of Science, Kyushu University, 744 Motooka, Nishi-ku, Fukuoka 819-0395, Japan; Department of Biosciences and Informatics, Faculty of Science and Technology, Keio University, 3-14-1, Hiyoshi, Kohoku-ku, Yokohama, Kanagawa 223-8522, Japan

## Abstract

Coherent Raman imaging (CRI) enables label-free chemical imaging based on intrinsic molecular vibrations, but its applicability is often limited by low sensitivity, hindering the detection of low-abundance biomolecules. While electronic resonance significantly enhances sensitivity, most resonance CRI implementations rely on narrowband excitation and/or detection, which limits spectral coverage and makes it difficult to distinguish target molecules from backgrounds. Here we address these challenges by developing broadband electronic-resonance coherent anti-Stokes/Stokes Raman scattering (BER-CARS/CSRS) microscopy. We show that BER-CARS/CSRS enables highly sensitive, label-free imaging of endogenous chromophores with broad spectral coverage of the entire fingerprint region. Specifically, we captured time-lapse images across the fingerprint region, visualizing low-abundance cytochromes alongside abundant biomolecules (lipids, proteins, nucleic acids) in living HEK293 cells. Furthermore, we applied the method to complex biological tissues, mapping distinct distributions of cytochromes in mouse brain slices, highlighting their characteristic localizations in the cortex and the cerebral ventricle wall. Our results demonstrate that BER-CARS/CSRS is a powerful platform for highly sensitive chemical imaging, from large-area tissue mapping to organelle-level dynamics. By coupling resonance enhancement with broadband fingerprinting, BER-CARS/CSRS enables dynamic, label-free phenotyping of mitochondrial and physiological states from single cells to tissues, opening a path to quantitative, slide-scale chemical histopathology and intraoperative margin assessment.

## Introduction

Coherent Raman imaging (CRI) is a powerful optical modality that enables label-free visualization of biomolecules in living cells and tissues by detecting their intrinsic vibrational frequencies. In CRI, molecular vibrations are coherently excited by nonlinear interactions induced by two-color laser fields, producing coherent optical signals that can be recorded with high spatial and temporal resolution. Representative implementations include coherent anti-Stokes Raman scattering (CARS) and stimulated Raman scattering (SRS) ^1-5^, which offer rapid image acquisition, three-dimensional sectioning capability, and molecular specificity without the need for fluorescent labels. These techniques have been applied across diverse contexts, such as *in vivo* monitoring of lipid droplets^6,7^, nucleic acids^2,8-10^, *ex vivo* visualization of amyloid plaques^11^ and brain tumors^12^, identification of microplastics in zooplankton^13^, label-free histopathological diagnosis of metabolic diseases^14^, gene-correlated metabolic imaging^15^, and *in vivo* tracking of drug delivery carriers^16^. Despite these advantages, the sensitivity of CRI techniques is typically limited to the millimolar range due to the inherently small Raman scattering cross section, making it challenging to visualize low-abundance molecules.

To overcome the sensitivity limit in CRI, harnessing electronic resonance is a promising approach. In electronic resonance Raman processes, the excitation wavelength is tuned within the absorption band of molecules, resulting in a dramatic increase in the Raman scattering cross section for vibrational modes coupled to the electronic transition^17-19^. In spontaneous Raman imaging, resonance Raman enhancement has widely been utilized to visualize light-absorbing biomolecules such as carotenoids and cytochromes^20-22^. While electronic resonance CRI^23-26^ has proven to be a promising approach due to its high sensitivity down to sub-micromolar level, its full potential has not yet been unlocked because most implementations rely on narrowband excitation and detection. Specifically, narrowband-detection CRI has two major drawbacks. First, molecular fingerprinting ability of Raman measurements is compromised, resulting in ambiguous interpretation of the obtained Raman images. Second, in coherent Raman processes in particular, vibrationally resonant signals are frequently obscured by non-specific background from four-wave mixing or cross-phase modulation. In these cases, broadband detection is crucial for resolving overlapping peaks and enabling quantitative interpretation of the spectra.

In this work, we address these limitations by developing a broadband electronic-resonance coherent anti-Stokes and Stokes Raman scattering (BER-CARS/CSRS) microscopy platform. Our system employs a near-infrared (NIR) supercontinuum (SC) source as the pump and Stokes fields, combined with a narrowband visible laser as the probe field, enabling broadband spectral acquisition across the fingerprint region under strong electronic resonance conditions. This configuration integrates the high sensitivity of electronic resonance with the wide spectral coverage needed for comprehensive molecular fingerprinting. Using BER-CARS/CSRS microscopy, we achieved label-free detection and imaging of endogenous light-absorbing biomolecules, including cytochromes in living HEK293 cells and mouse brain slices. These results highlight the capability of our method to provide quantitative, chemically specific imaging for a broad range of applications in life sciences and biomedical research.

## Results and Discussion

CARS and CSRS are third-order nonlinear optical processes induced by the pump, Stokes, and probe fields (Fig. 1a). In these processes, the pump and Stokes fields create vibrational coherences of the molecules, which subsequently interact with the probe field to generate a third-order nonlinear polarization that radiates CARS or CSRS fields. In electronic-resonance CARS or CSRS (ER-CARS, ER-CSRS), the photon energy is tuned to match an electronic transition energy of the molecule, resulting in resonance enhancement of the third-order nonlinear susceptibility χ^(3)^ and a significant enhancement in signal intensity (Fig. 1a). To enable both broadband spectral acquisition and visible electronic enhancement, BER-CARS and BER-CSRS employ broadband pump and Stokes pulses in the NIR region (900 -1300 nm) and a narrowband probe pulse in the visible region (520 nm) (Fig. 1b). The schematic of our developed BER-CARS/CSRS imaging system is shown in Fig. 1c (see the Methods section for details). All the pulses were derived from a single femtosecond laser. To obtain the broadband pump (*ω*_1_) and Stokes (*ω*_2_) pulses, a portion of the laser output was coupled into a photonic crystal fiber (PCF) to generate an NIR femtosecond SC spanning from 900 to 1300 nm and subsequently compressed with a chirped mirror pair and a prism compressor. The narrowband probe (*ω*_3_) pulse was obtained by second harmonic generation using a nonlinear optical medium and wavelength selected with a 4-f filter. The two pulses were temporally and spatially overlapped using an optical delay line and a wavelength filter and guided into a custom-built microscope. The generated CARS and CSRS signals were spectrally filtered with a notch filter and a short-pass filter and detected using a home-built spectrometer and a CCD camera. Figure 1d shows a BER-CARS/CSRS spectrum obtained from liquid cyclohexane in 1-ms exposure time, showing symmetric spectral profiles on both the Stokes and anti-Stokes sides and vibrational peaks consistent with reported Raman spectra of cyclohexane. The input power dependence of the CARS/CSRS signal intensities (Figs. 1e, 1f) shows a quadratic dependence on the pump/Stokes (NIR broadband) power and a linear dependence on the probe (visible narrowband) power. These dependences are consistent with the assumption that the CARS/CSRS signals were generated in the optical processes shown in Fig. 1b.

**Fig. 1.**
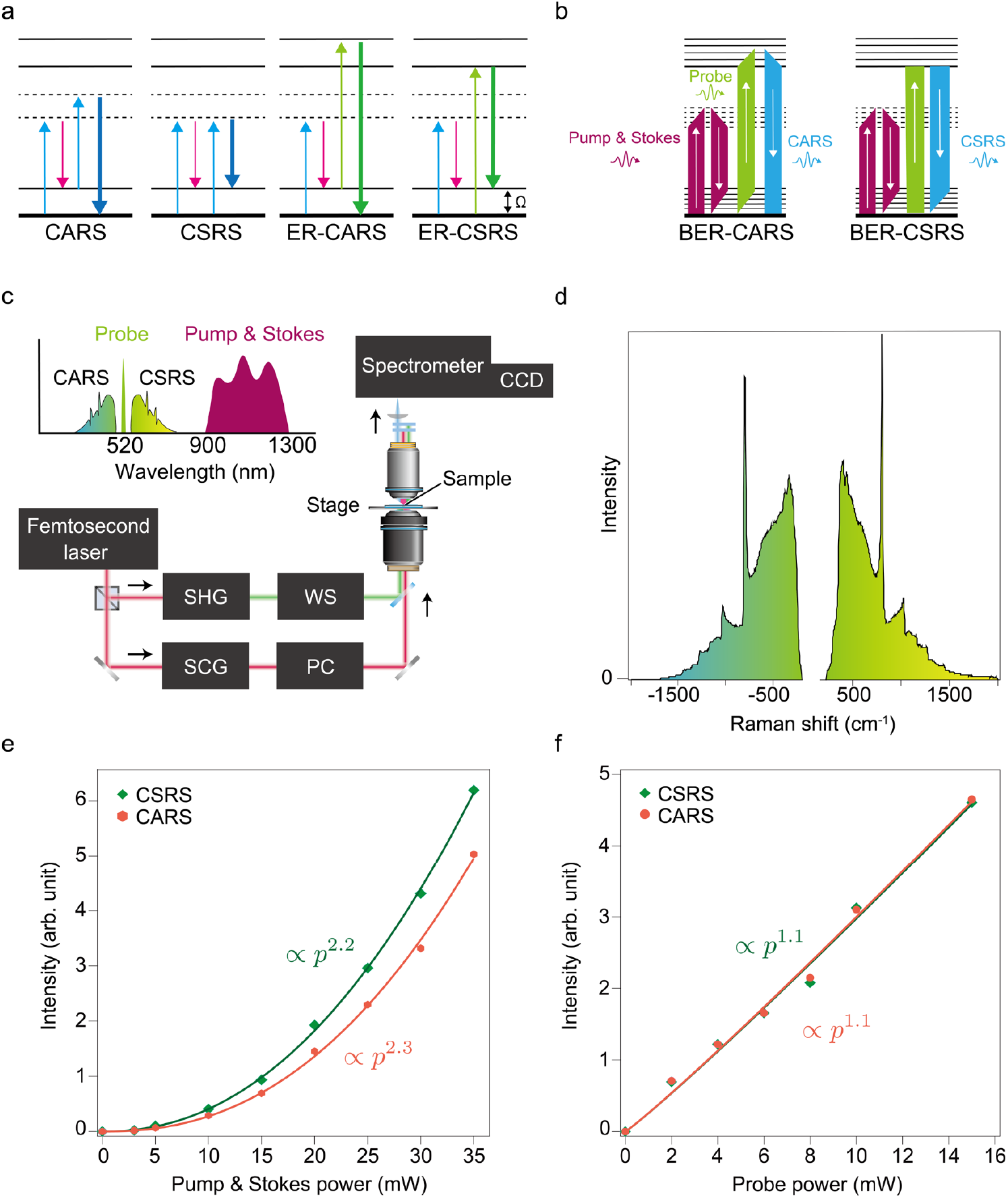
Concept and working principle of broadband electronic resonance CARS/CSRS (BER-CARS/CSRS) microscopy. **(a)** Energy diagrams of CARS/CSRS and ER-CARS/CSRS processes. **(b)** Energy diagram of BER-CARS/CSRS process. **(c)** Schematic of our BER-CARS/CSRS setup. Inset shows spectral profiles of the incident fields and the generated CARS and CSRS fields. SHG: Second harmonic generation, WS: wavelength selector, SCG: supercontinuum generation, PC: pulse compressor. **(d)** Raw BER-CARS/CSRS spectrum of liquid cyclohexane. **(e)** Power dependence of BER-CARS/CSRS signals on the pump and Stokes intensity. **(f)** Power dependence of BER-CARS/CSRS signals on the probe intensity.

To evaluate the basic performance of our BER-CARS/CSRS imaging system, we first performed the concentration-dependence study with water solutions of dimethyl sulfoxide (DMSO) and acetone solutions of astaxanthin at various concentrations. In this study, we focused on the CARS side of the spectra because clearer vibrational peaks of astaxanthin were detected in CARS compared to in CSRS due to lower fluorescence hindrance and higher resonance enhancement in the anti-Stokes region (*λ*_max_ = 477 nm for astaxanthin). Figures 2a, 2b show CARS spectra of DMSO-water solutions, where the distinct peak at 670 cm^-1^ is clearly visible at 40 mM (10 ms) and 5 mM (1000 ms). Figures 2c, 2d show CARS spectra of astaxanthin-acetone solutions, where the distinct peak at 1152 cm^-1^ is clearly visible at 10 μM in both exposure times. Due to the resonance enhancement, comparable signal intensities were obtained from astaxanthin at concentrations 3-4 orders of magnitude lower than those required for DMSO under the same exposure time and the excitation power. The limit of detection in BER-CARS in 10-ms exposure, which is a typical pixel dwell time in CARS imaging, is roughly 40 mM for DMSO (without electronic resonance) and 10 μM for astaxanthin (with electronic resonance). These results show that the resonance enhancement under visible excitation significantly improves the detection limits of label-free imaging of visible-light-absorbing biomolecules.

**Fig. 2.**
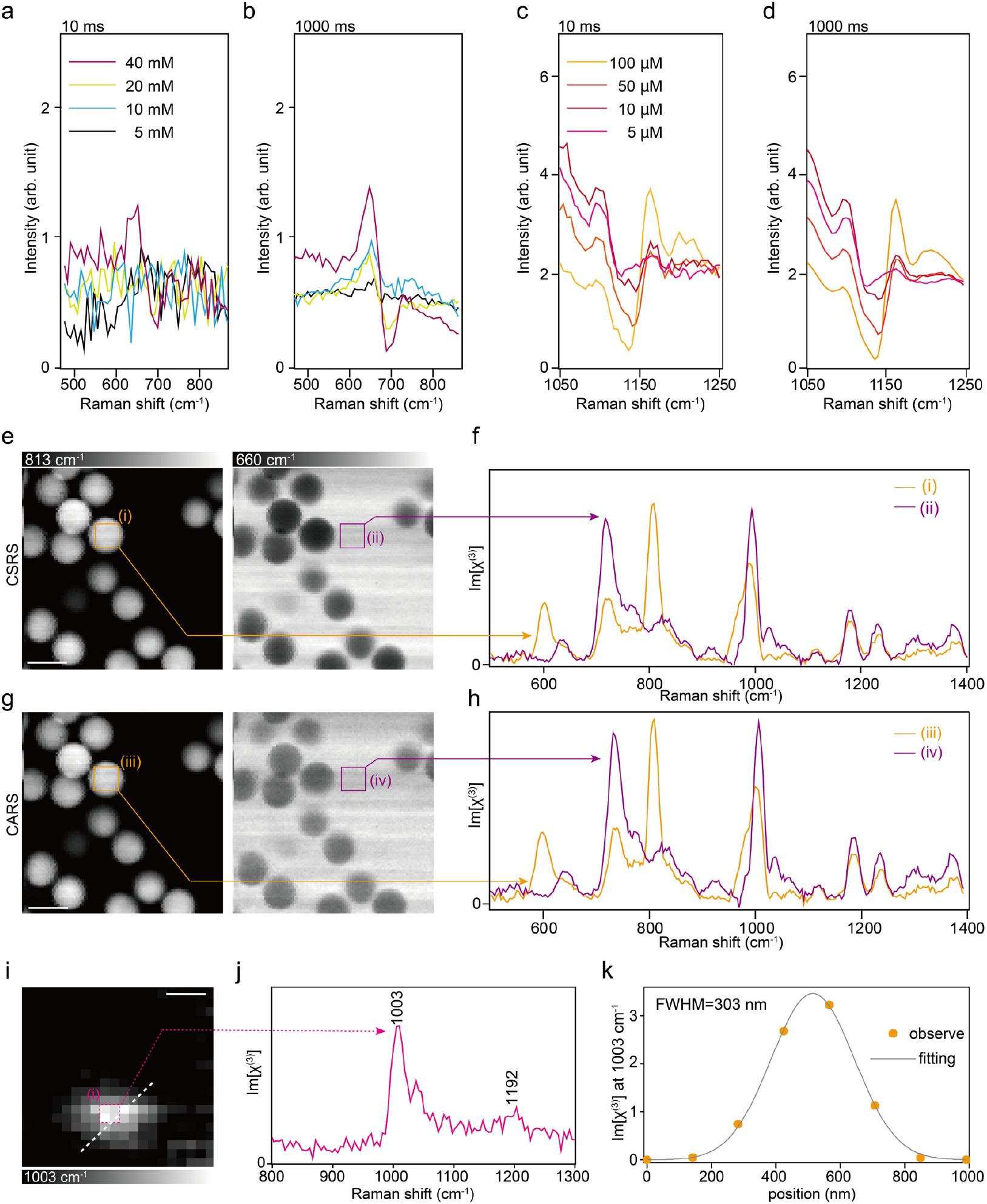
Concentration dependence and imaging performance characterization. **a, b)**CARS spectra of DMSO solutions at concentrations of 40, 20, 10, and 5 mM at exposure times of **(a)** 10 ms and **(b)** 1000 ms. **(c, d)** CARS spectra of astaxanthin-acetone solutions at concentrations of 100, 50, 10, and 5 μM at exposure times of **(c)** 10 ms and **(d)** 1000 ms. **(e)** BER-CSRS images of PMMA beads at 813 and 660 cm^−1^. **(f)** Average Im[χ^(3)^] spectra from regions (i) and (ii) in (**c)**. (**g**) BER-CARS images of PMMA beads at 813 and 660 cm^−1^. **(h)** Average Im[χ^(3)^] spectra from regions (iii) and (iv) in **(e)**. The image size: 101 × 101 pixels, pixel size: 0.5 μm x 0.5 μm, exposure time: 20 ms/pixel, Scale bar: 10 μm. (**i)** BER-CARS image of a 200 nm polystyrene bead at 1003 cm^−1^. The image size: 20 × 20 pixels, pixel size: 100 nm x 100 nm, exposure time: 50 ms/pixel, Scale bar: 400 nm. **(j)** Average Im[χ^(3)^] spectra from the boxed region in **(g). (k)** Intensity profile along the dashed line in **(i)**. Orange dots indicate observed values, and the gray curve indicates Gaussian fitting.

We next evaluated the imaging performance of our setup using standard test samples. Polymethyl methacrylate (PMMA) beads with a diameter of 10 μm were suspended in immersion oil (Immoil-F30CC, Olympus, Japan) and imaged by raster scanning the sample using a translational stage (Figs. 2e-2h). For quantitative interpretation of the CARS/CSRS spectra, we extracted the imaginary part of χ^(3)^ by using the maximum entropy method^27^. Image reconstruction was performed based on the magnitude of Im[χ^(3)^] at 810 cm^−1^ and 660 cm^−1^, where representative peaks of PMMA and the immersion oil are present, respectively. The CARS (Fig. 2e) and CSRS (Fig. 2g) exhibited similar profiles, which indicates that CARS and CSRS provide identical information in the non-resonant case. Figures 2f, 2h show average Im[χ^(3)^] spectra derived from CARS and CSRS measured in the regions identified by the rectangles in the images. The PMMA bead and the immersion medium are spectrally distinguished with rich chemical information in the entire fingerprint region. It is worth mentioning that the intensity ratio of CARS to CSRS differs depending on the sample thickness in the epi-detection configuration^28^, making CARS/CSRS dual measurement a promising tool for sensing the sample thickness.

We further assessed the capability of imaging nanoscale samples by imaging a 200-nm polystyrene (PS) bead. Figure 2i shows a CARS image consisting of Im[χ^(3)^] at 1003 cm^−1^, where a characteristic Raman peak of PS exists (Fig. 2j). This image indicates that our imaging platform clearly resolves submicron structures. The Im[χ^(3)^] profile across a bead is shown in Fig. 2j, where Gaussian fitting analysis reveals that the full width at half maximum (FWHM) of the peak is approximately *d*_obs_ = 303 nm (Fig. 2k). We estimated the FWHM of the point spread function by 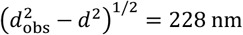, where *d* = 200 nm is the diameter of the bead. This value is larger than the theoretical estimation of 130 nm, estimated based on the diffraction limit of the 520-nm and 1040-nm beams with third-order nonlinear interaction and an objective numerical aperture of 1.42. We think this mismatch comes from imperfect spatial overlaps of the excitation beams, especially along the axial direction and non-ideal beam profile of the output of the PCF.

We applied BER-CARS/CSRS imaging to live HEK293 cells, a widely used human-derived cell line that serves as a convenient model for investigating fundamental cellular processes, including mitochondrial function^29,30^. Cytochromes in these cells are central to energy metabolism^31^ and apoptosis^32^, yet their spatiotemporal dynamics, such as rapid redox changes and redistribution under stress, remain incompletely understood due to limited temporal resolution of conventional Raman approaches. Demonstrating that BER-CARS/CSRS can directly detect these chromophores in living cells thus represents a critical step toward elucidating their dynamic roles in cellular metabolism. Figure 3a shows BER-CSRS images of living HEK293 cells reconstructed by Im[χ^(3)^] at 1443, 795, 750 and 1001 cm^−1^, where the vibrational peaks of CH_2_ scissoring, pyrimidine ring/PO_2_^?^ backbone vibrations, heme breathing modes and phenylalanine ring breathing, are found, respectively. The spatial profiles in the 1443-cm^-1^ and 1001-cm^-1^ images primarily reflect overall cell morphology, consistent with previously reported Raman images of lipids and proteins. In addition, the 795-cm^−1^ image exhibited strong intensity in the nuclei and nucleoli, indicating that the signal at this frequency reflects the local abundance of nucleic acids. The image at 750 cm^−1^, where no clear contrast has previously been reported in conventional CRI bioimaging, shows high intensity at peripherals of the nuclei. Considering the reported Raman peaks of cytochromes at 750 cm^-1^ and localization of cytochromes around the nuclei^20,33^, these results suggest that our BER-CSRS platform successfully visualizes cytochromes in living cells. Figure 3b presents averaged Im[χ^(3)^] spectra from three regions indicated in inset, enriched in nucleic acids (i), cytochromes (ii) and proteins (iii). Region (i) shows prominent peaks at 795 cm^−1^, (ii) at 750 cm^−1^, and (iii) at 1001 cm^−1^, further supporting the above assignment of the signals.

**Fig. 3.**
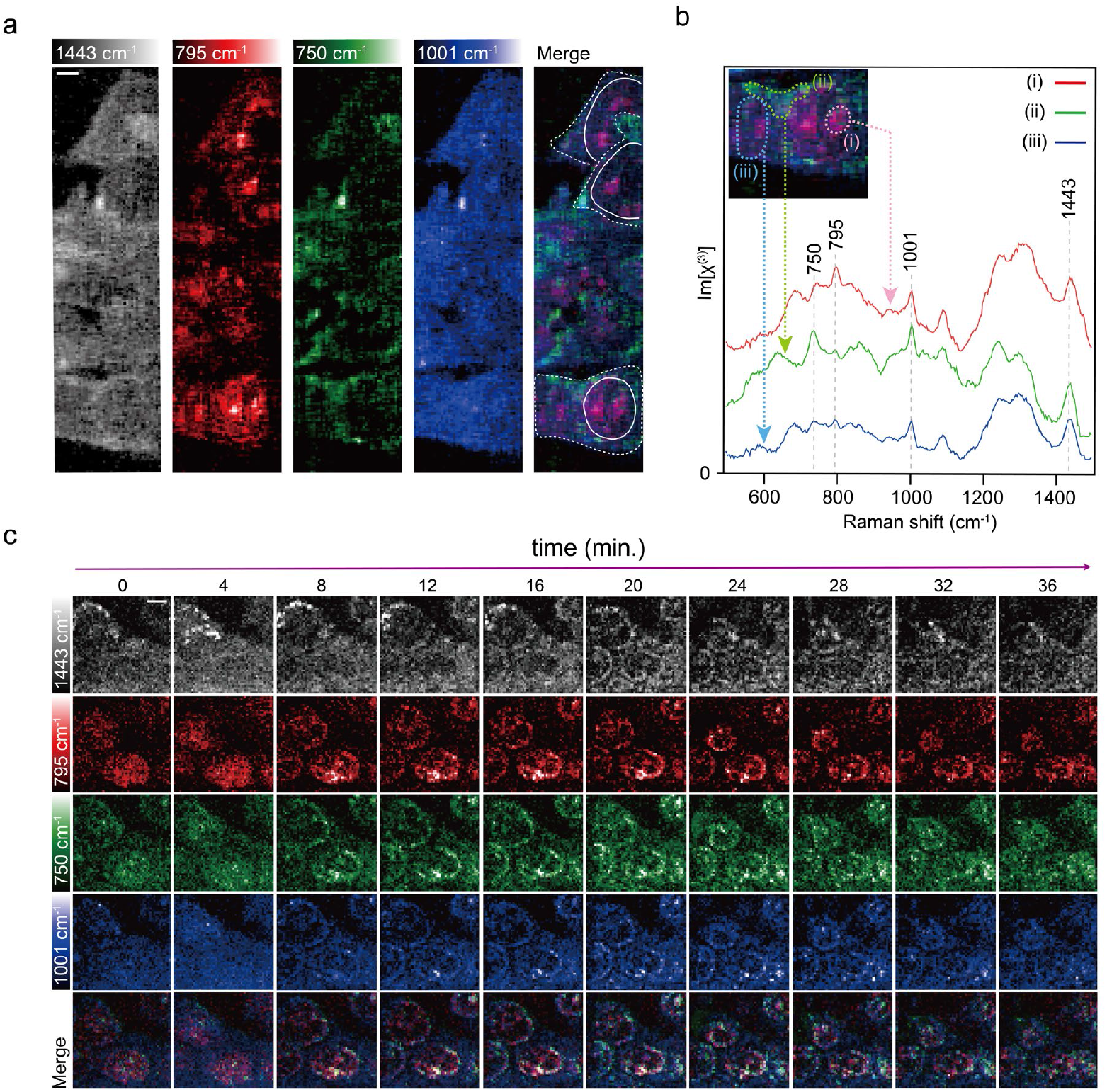
BER-CSRS imaging of living HEK293 cells. **(a)** BER-CSRS images of living HEK293 cells reconstructed at 1443 (lipids), 795 (nucleic acids), 750 (cytochromes) and 1001 cm^−1^ (proteins). Solid white line: nucleus; dashed white line: cell boundary. Image size: 201 × 45 pixels, pixel size: 0.5 μm x 0.5 μm, exposure time: 20 ms/pixel. Scale bar: 5 μm. **(b)** Average Im[χ^(3)^] spectra corresponding to regions (i)–(iii) in inset. The inset shows a magnified view of the cell in (a).**(c)** Time-lapse BER-CSRS imaging of HEK293 cells during apoptosis at 1443 (lipids), 795 (nucleic acids), 750 (cytochromes) and 1001 cm^−1^ (proteins). BER-CSRS imaging was initiated at *t* = 0 min and performed until *t* = 36 min with a 4-min interval. The image size: 51 × 51 pixels, pixel size: 0.5 μm x 0.5 μm, exposure time: 1 ms/pixel, Scale bar: 5 μm.

We next performed time-lapse BER-CARS/CSRS imaging of HEK293 cells undergoing apoptosis (Figs. 3c, Supplementary Movie 1). HEK293 cells were incubated in Hanks’ balanced salt solution (HBSS) for 1 h, mounted on a slide, with the observation starting at *t* = 0 min. The reconstructed BER-CSRS images captured dynamic changes in lipids (1443 cm^−1^), nucleic acids (795 cm^−1^), cytochromes (750 cm^−1^), and proteins (1001 cm^−1^) in a cell. In the early stage, cytochromes were uniformly distributed within the nucleus (Fig. 3c, *t* = 0 − 4 min). This is consistent with early apoptotic nuclear translocation of cytochrome c, implicated in chromatin condensation and caspase activation^34^. Actually, BER-CSRS image at 795 cm−^1^ taken at *t* = 8 min (Fig. 3c) shows spatial patterns indicative of chromatin condensation. Remarkably, nuclear cytochromes colocalize with nucleic acids inside the nucleus during apoptosis (Fig. 3c). These observations are consistent with the models in which cytochromes regulate chromatin during the DNA-damage response^34,35^ and, to our knowledge, provide the first label-free confirmation of this sequence without fluorescent probes.

To further showcase the utility of BER-CARS for visualizing complex tissue, we imaged mouse brain slices and leveraged electronic-resonance enhancement to detect a cytochrome-associated heme band near 750 cm−^1^. Because mitochondria govern neuronal energy supply and signaling, and because their remodeling modulates brain-state control, mapping of mitochondrial distribution provides biologically important insights^36-38^. First, we acquired a large-area scan spanning the cerebral cortex, white matter, and hippocampus (Figs. 4a - region (i)). These macroscopic regions are readily distinguished by chemical contrast: the nucleic-acid band at 790 cm−^1^ is enriched in cell-body–dense cortical layers and the CA1 region, whereas the 1005 cm−^1^ band—attributed to proteins, appeared relatively uniform across the brain tissue. Also, cytochromes are visualized by image reconstruction at 750 cm−^1^, showing non-localized distribution of cytochromes in cytoplasm. Broadband and electronic-resonance capability of the present method enables multimodal chemical imaging including low-abundance molecules like cytochromes.

**Fig. 4.**
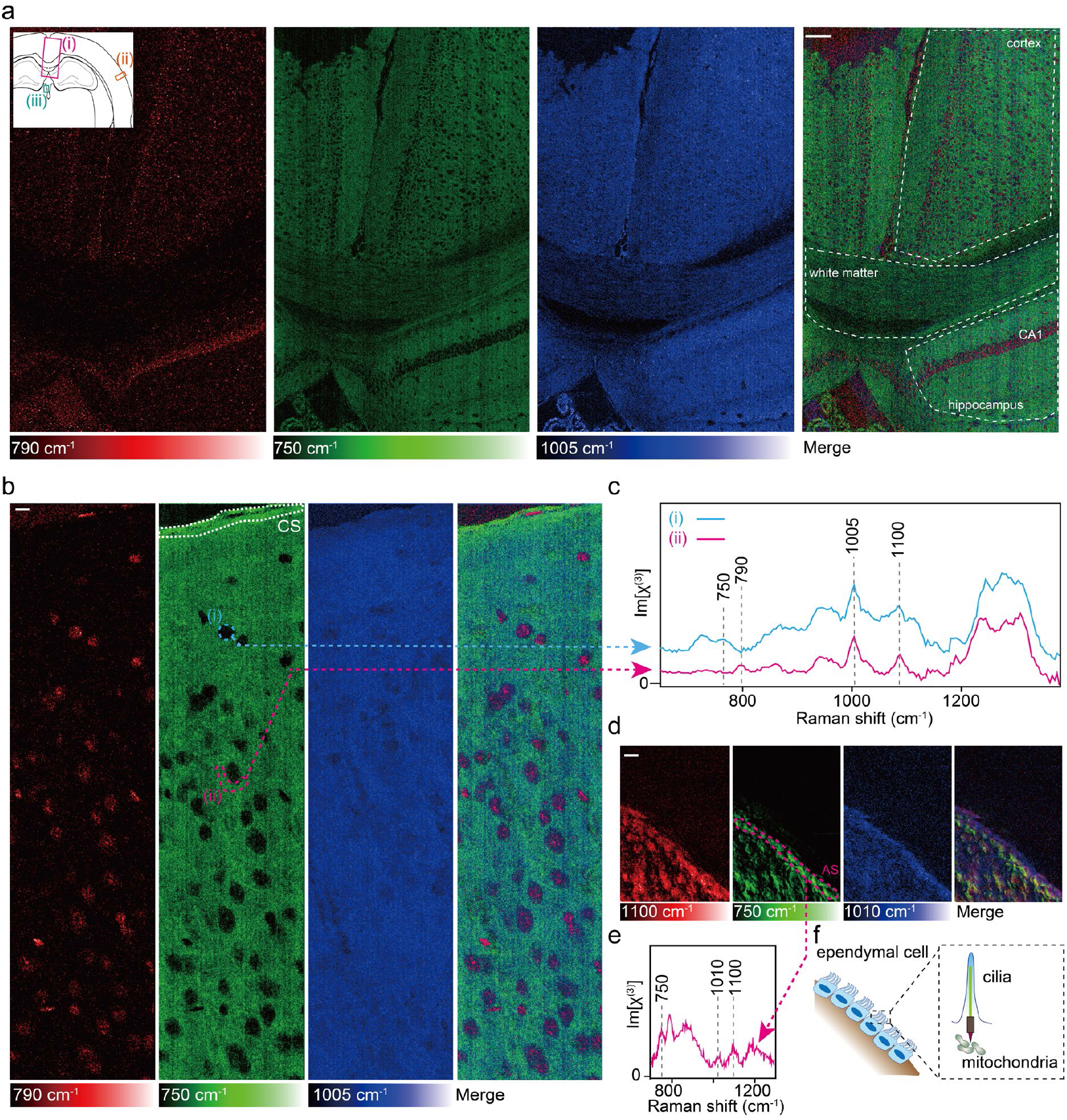
Chemical imaging of mouse brain tissue by BER-CARS microscopy. **(a)** Large-area BER-CARS imaging of mouse brain across cortex, white matter, and hippocampus. The region indicated as (i) in the inset was measured. Signals at 790, 750, and 1005 cm^−1^ correspond to nucleic acids, cytochromes, and proteins, respectively. Image size: 801 × 501 pixels, pixel size: 2 μm x 2 μm, exposure time: 5 ms/pixel. Scale bar: 100 μm. **(b)** High-resolution BER-CARS imaging of the cortex. The area measured at (ii) in the inset of **(a)**. CS: cortical surface. Image size: 801 × 189 pixels, pixel size: 0.5 μm x 0.5 μm, exposure time: 5 ms. Scale bar: 5 μm. **(c)** Average Im[χ^(3)^] spectra corresponding to regions (i) and (ii) in **(b). (d)** High-resolution BER-CARS imaging around the cerebral ventricle wall. The area measured at (iii) in the inset of **(a)**. AS: apical surface. Signals at 1100, 750, and 1010 cm^−1^ correspond to nucleic acids, cytochromes, and proteins, respectively. Image size: 101 × 161 pixels, pixel size: 0.5 μm x 0.5 μm, exposure time: 5 ms/pixel. Scale bar: 10 μm. **(e)** Average Im[χ^(3)^] extracted from the region indicated by the magenta dotted line in **(d). (f)** Schematic illustration of the measurement region. Mitochondria are localized at the base of the cilia in ependymal cells.

We next performed higher-resolution imaging of the cortex (Figs. 4a - region (ii), 4b, 4c), resolving chemical features at the single-cell level. The 790 cm−^1^ signal localized to nuclei, sharply delineated from the cytoplasm, while the ∼750 cm−^1^ cytochrome-associated signal was broadly detected in cytoplasm and the1005 cm−^1^ proteins Notably, the mean intensity of the ∼750 cm−^1^ band is higher near the cortical surface (CS). A similar mitochondrial behavior has been observed in fluorescence image of mouse brain, where HK1 was stained and higher fluorescence intensity was detected at the brain surfaces^39,40^. Explicit discussion about the higher mitochondrial concentration at the surface was not provided in this work presumably because higher fluorescence signals at the surface can be an artifact during immunostaining due to higher exposure to the fluorescence dye at the tissue peripheral. As CARS directly detects cytochromes without labeling, the higher mitochondrial concentration at the surface is now more evident. While the underlying biological mechanisms need to be further investigated in future study, it may be related to high metabolic activity at the axon terminal^41,42^, which necessitates mitochondria.

Finally, we performed BER-CARS imaging of the surrounding cerebral ventricle wall (Figs. 4a - region (iii), 4d-4f). Ependymal cells, lining the cerebral ventricle wall, are known to possess cilia for circulating cerebrospinal fluid, and their active motility is energetically supported by numerous mitochondria densely packed at the ciliary base^43,44^. Figure 4d shows BER-CARS images of ependymal cells at 1100, 750, and 1010 cm−^1^, which correspond to the vibrational peaks of PO_2_− stretching, heme breathing modes, and phenylalanine ring breathing, respectively. These peaks are attributed to nucleic acids, cytochromes, and proteins. In the BER-CARS image, we found localized cytochrome signals on the apical surface. The ability of our method to visualize mitochondrial cytochromes in ependymal cells can be an important tool to study ciliopathy because the lack of mitochondrial activity is one important origin of ciliopathies, such as diabetes, infertility, and obesity^45^. These results demonstrate that BER-CARS microscopy is an innovative imaging platform that non-invasively provides multi-scale chemical information of biological tissues, ranging from large-area tissue mapping and subcellular molecular distributions to the visualization of functional organelles.

## Conclusion

We developed a BER-CARS/CSRS microscope that brings electronic-resonance enhancement to coherent Raman imaging while retaining broadband vibrational fingerprinting. By extracting the imaginary component of χ^(3)^, the platform provides quantitatively interpretable, background-mitigated spectra across the fingerprint region. These capabilities enabled label-free, time-lapse visualization of endogenous cytochromes (e.g., cytochrome *c*) in living cells and robust mapping of their distributions in complex mouse brain tissue. In particular, we observed spatial patterns ranging from mitochondria-associated structures to enriched signals at the apical surface of ependymal cells, consistent with known mitochondrial accumulation at this interface.

Besides the biological applications shown in the present work, our BER-CARS/CSRS imaging platform has the potential to elucidate various biological phenomena in cells and tissues. The combination of broadband coverage and resonance enhancement is well suited to tracking redox-dependent spectral changes of cytochromes, informing analyses of mitochondrial electron-transport activity, apoptotic signaling, and cellular responses to oxidative stress, as well as activity-dependent metabolism in neural circuits. The same principles extend to other key endogenous chromophores, including flavins, melanin, and hemoglobin, and can be leveraged to push the sensitivity of Raman tags toward highly specific, multiplexed molecular tracking.

While the present BER-CARS/CSRS microscope is already a powerful tool in biological research, its performance can further be enhanced in multiple directions. First, implementing epi-detection for both CARS and CSRS will enable size-selective measurements as epi-detected CARS and CSRS show different size-dependence. Simultaneous forward/epi readout across CARS and CSRS therefore provides complementary sensitivity that helps resolve subwavelength structure. Second, by generating broadband pump/Stokes light in the visible with sufficient difference-frequency coverage (∼580 - 670 nm), the system can interrogate the CH-stretch region (∼2800 - 3100 cm^−1^) concurrently with the fingerprint band, enabling imaging with richer chemical specificity. Third, co-registration with other nonlinear contrasts, including SHG, THG, and two-photon fluorescence, will yield a higher multimodality that fuses structural and morphological information with chemically specific BER-CARS/CSRS maps. Finally, integration with information science (task-specific priors, physics-informed spectral unmixing, active acquisition, and anomaly detection) can drive more goal-oriented measurements, for instance, pathology triage, cell-cycle staging, or discovery of atypical regions, reducing dwell time and dose while increasing diagnostic yield.

## Methods

### Experimental Setup for BER-CARS/CSRS Microscopy

Our home-built BER-CARS/CSRS microscopy setup is illustrated in Fig. S1. The light source was a femtosecond laser (FLINT FL2-SP, Light Conversion, Lithuania) operating at 1040 nm with a 50-fs pulse duration and a 76-MHz repetition rate. The output was split into two arms to generate near-infrared pump–Stokes pulses and a visible probe pulse. For the pump and Stokes beams, a fraction of the output was coupled into a photonic crystal fiber (LMA-PM-5, Thorlabs, Japan) to produce a supercontinuum. An automatic fiber-alignment controller (KNA-VIS, Thorlabs, Japan) mounted on a three-axis stage (MAX313D, Thorlabs, Japan) was used for stable coupling and wavelength tuning. Group-delay dispersion introduced by the optics was compensated with chirped mirrors and a prism-pair compressor. For the probe beam, the remaining output was frequency-doubled in a periodically poled lithium niobate (PPLN) crystal to 520 nm by second-harmonic generation (SHG). The SHG beam was spectrally narrowed with a 4-f spectral filter (grating–lens-slit–lens-grating), yielding a narrowband probe. A motorized delay line in the probe arm provided precise temporal overlap with the pump–Stokes pulses at the sample plane. The pump, Stokes, and probe beams were combined collinearly with a dichroic mirror and delivered to a laser-scanning microscope. The beams were focused onto the sample with an oil-immersion objective (UPLSAPO 60×/NA 1.42, Olympus, Japan), and the generated CARS and CSRS signals were collected with a second objective (Plan Apo 20×/NA 0.75, Nikon, Japan). The sample was mounted on a motorized linear stage (MLS203-2, Thorlabs, Japan) for scanning. Excitation light was blocked with appropriate notch and long-pass filters, and the signals were dispersed by a custom-made spectrometer and detected with a charge-coupled device (CCD) camera (PIXIS 100BR-DD eXcelon, Princeton Instruments, USA). The imaginary part of χ^(3)^, which is equivalent to the spontaneous Raman spectrum, was retrieved from the raw CARS/CSRS data using the maximum entropy method^27^. Im[χ^(3)^] spectra were preprocessed using the rolling-ball algorithm for background subtraction. Imaging data of HEK293 cells were analyzed by singular value decomposition (SVD) (Fig. 3a) and Tucker decomposition (Fig. 3c), where a difference Raman intensity at two wavenumbers was mapped. For brain tissue (Fig. 4), intensity mapping was performed using amplitudes obtained from Gaussian fitting.

### Standard samples preparation

Polymethyl methacrylate (PMMA) beads (10 µm diameter; Polysciences Inc., USA) were suspended in immersion oil (Immoil-F30CC, Olympus, Japan), and polystyrene beads (200 nm diameter; Polysciences Inc., USA) were dispersed in water. Each sample was mounted between a coverslip and a microscope slide and sealed with nail polish.

### Cell culture

HEK293 cells (RIKEN BRC Cell Bank) were cultured on coverslips in Dulbecco’s modified Eagle’s medium (DMEM; Fujifilm Wako, Japan) supplemented with 10% fetal bovine serum at 37 °C in a humidified 5% CO_2_ atmosphere for 3 days. Before imaging, coverslips were rinsed in Hanks’ balanced salt solution (HBSS) and incubated for 1 h. The coverslip with adherent cells was then inverted onto a glass slide and sealed with nail polish.

### Animals and brain tissue preparation

C57BL/6N wild-type male mice were housed under controlled temperature and humidity on a 12-h light/12-h dark cycle. Mice were deeply anesthetized with 5% isoflurane followed by intraperitoneal somnopentyl and transcardially perfused with phosphate-buffered saline (PBS), followed by 4% paraformaldehyde (PFA) in PBS. The brains were dissected and post-fixed in 4% PFA in PBS at 4 °C overnight, cryoprotected sequentially in 15% sucrose for 1 day and 30% sucrose for 1 day, embedded in OCT compound (Sakura Finetek), and frozen at −80 °C. The brains were cryosectioned coronally at 40 µm thickness. Sections were rinsed and mounted with PBS. All animal experimental procedures were approved (Approved protocol ID # 24-324) and conducted following the guidelines established by the Institutional Animal Care and Use Committee of the University of Tsukuba. In the imaging experiments, data shown in Fig. 4a–c were obtained from the same brain, whereas those in Fig. 4d and 4e were obtained from different mice.

## Acknowledgements

The authors thank W. Yamamoto, T. Sasaki, R. Harada and M. Mashima for their helpful discussion and technical support. This research was supported by JST FOREST (JP21470594 to K.H.), JSPS KAKENHI Grant-in-Aid for Scientific Research (B) (JP23K23297 to K.H., JP25K03136 to K.H., JP24K03251 to H.K.), JSPS KAKENHI Grant-in-Aid for Scientific Research (A) (JP21H04961 to H.K.), Grant-in-Aid for Challenging Exploratory Research (JP23K17933 to H.K.), and JST Mirai Program (JPMJMI22G5 to H.K.), Grant-in-Aid for Transformative Research Areas (JP25H01396 to K.H., JP25H01393 to K.H.), Grant-in-Aid for Challenging Research (Pioneering) (JP25K21710 to K.H.), Grant-in-Aid for Scientific Research (S) (JP25H00410 to K.H.), Grant-in-Aid for JSPS Fellows (JP23KJ0267 to Y.M, JP23KJ0286 to M.M.), Grant-in-Aid for Early-Career Scientists (JP24K17623 to R.O.), Grant-in-Aid for Specially Promoted Research (JP22H04918 to M.Y.), Fund for the Promotion of Joint International Research (International Leading Research) (JP22K21351 to M.Y.), the Japan Agency for Medical Research and Development (AMED) (JP21zf0127005 to M.Y. and S.H.), the World Premier International Research Center Initiative (WPI) from the Ministry of Education, Culture, Sports, Science, and Technology (MEXT) (to M.Y.), Photographic Research Fund of Konica Minolta Imaging Science Foundation, and Murata Science and Education Foundation.

## Author contributions

K.H. conceived and directed the project. Y.M., M.M., R.O., and K.H. designed the study. Y.M. and K.H. performed the experiments. M.M. and M.Y. prepared the samples. Y.M., N.S. and K.H. analyzed the data. All authors wrote the paper.

## Competing interests

The authors declare no competing financial interests.

## Notes

### Competing Interest Statement

The authors have declared no competing interest.

