## Supplementary Figure 1 for "Broadband electronic resonance coherent anti-Stokes/Stokes Raman scattering microscopy"

[4]Department of Pharmacology, Keio University School of Medicine, 35 Shinanomachi, Shinjuku, Tokyo 160-8582, Japan

*Corresponding Authors

**Table of contents**

Supplementary Figure・・・・・・・・・・・・・・・・・・・・・・・・・・・・・・・S2

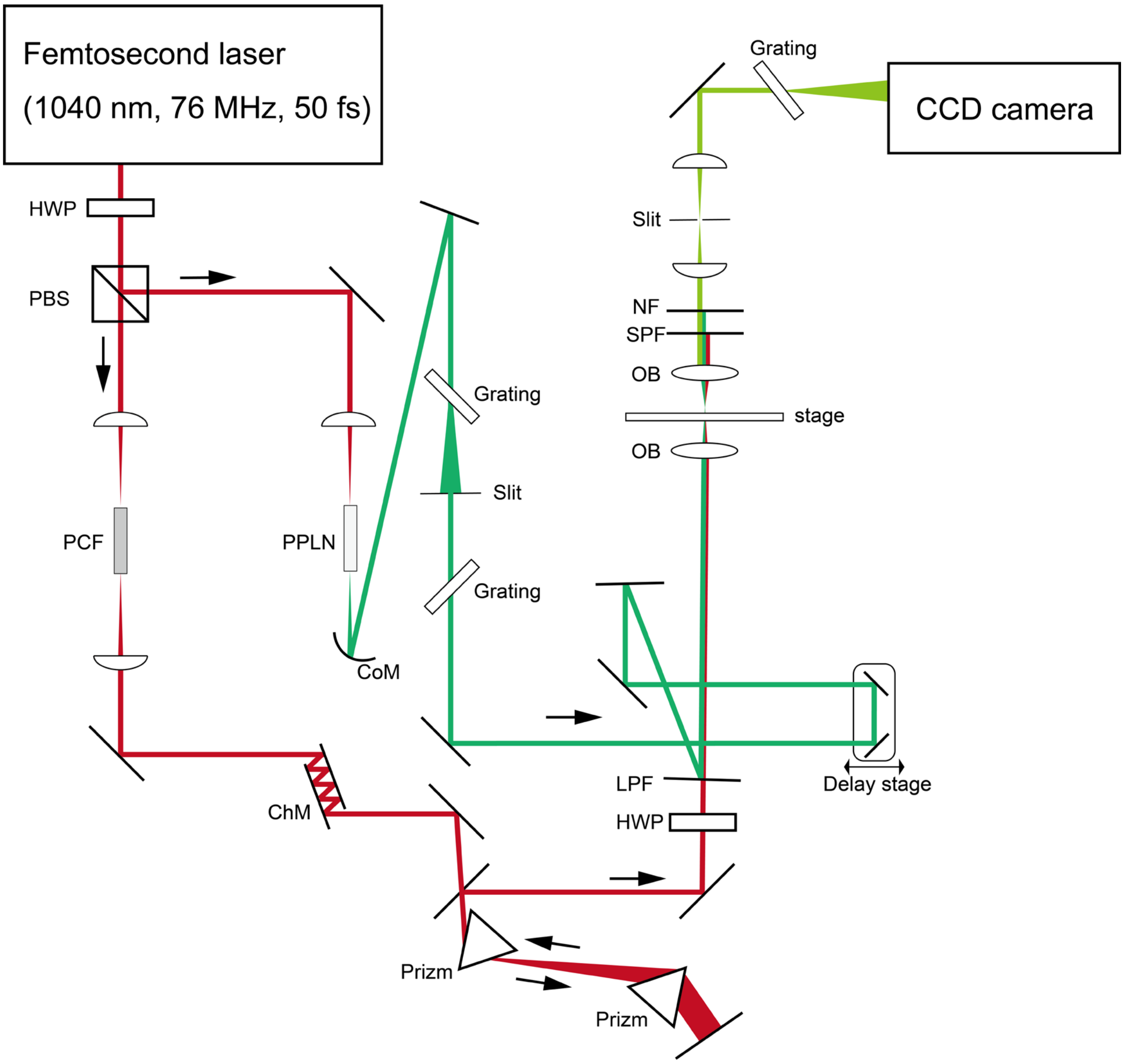


**Fig. S1**: Optical setup for BER-CARS/CSRS system. HWP: half-wave plate; PBS: polarizing beam splitter; PCF: photonic crystal fiber; PPLN: periodically poled lithium niobate; ChM: chirp mirror; CoM: concave mirror; LPF: long pass filter; OB: objective lens; SPF: short pass filter; NF: notch filter.
