## Supplementary figures and images for "Broadband electronic resonance coherent anti-Stokes/Stokes Raman scattering microscopy"

### Supplementary Movie 1

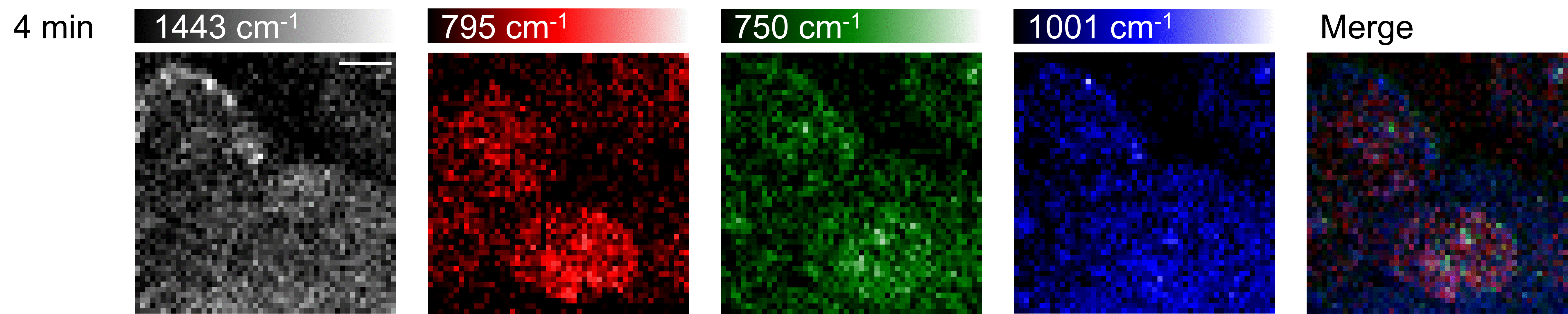
